# Proteins observed in scalp hair from preschool children

**DOI:** 10.1101/2021.01.22.427851

**Authors:** Kratika Singhal, Ryan D. Leib, Cynthia R. Rovnaghi, Maria Xenochristou, Nima Aghaeepour, Allis S. Chien, Grace K-Y. Tam, Monica O. Ruiz, Deendayal Dinakarpandian, Kanwaljeet J. S. Anand

## Abstract

Early childhood experiences have long-lasting effects on subsequent mental and physical health, education, and employment. Measurement of these effects relies on insensitive behavioral signs, subjective assessments by adult observers, neuroimaging and neurophysiological studies, or remote epidemiologic outcomes. Despite intensive search, no biomarkers for developmental changes in the brain have been identified. We analyzed scalp hair from healthy children and their mothers using an unbiased proteomics platform to reveal 1368 hair proteins commonly observed in children, 1438 proteins commonly observed in mothers, and 1288 proteins observed sporadically in individual subjects. Mothers showed higher numbers of peptide spectral matches and hair proteins compared to children, with important age-related differences between mothers and children. Age-related differences were also observed in children, with differential protein expression patterns between younger (2 years and below) and older children (3-5 years). Boolean analyses showed greater conservation of hair protein patterns between mothers and their biological children as compared to mothers and unrelated children. The top 5% proteins driving population variability represent biological pathways associated with brain development, immune signaling, and stress response regulation. Non-structural proteins observed in scalp hair may include promising biomarkers to investigate the developmental changes associated with early childhood experiences.

**One Sentence Summary:** The non-structural proteins observed in scalp hair from preschool children show evidence for heritability, reflect biological functions such as brain development, or immune function and regulation of stress responses, and exhibit age- and sex-related differences across periods of early childhood development.

## Introduction

Early human development remains exquisitely sensitive to parental, environmental, and societal influences that multiplex the history of each individual (via genetic and epigenetic factors) with their daily experiences. Variations in these factors, such as the social determinants of health, can singly or collectively introduce differences in their developmental outcomes(*1, 2*). Such differences are then magnified in the higher-order cognitive and behavioral capacities of humans, which are built on a series of sequential or staggered developmental epochs that can enable or constrain their future role(s) in society, as well as their mental and physical health(*3-5*).

Although the outcomes of early childhood are easily assessed, but *objectively* assessing their social, emotional, or other environmental inputs across multiple timescales is challenging. These challenges result from most subjects being pre-verbal, coming from unknown environments, or accompanied by unreliable historians(*1, 6*). Developmental timescales can also range from milliseconds to minutes (e.g., affecting acute neuromodulatory tone, neuronal oscillations, neuroendocrine changes), days to weeks (e.g., affecting circadian rhythms, metabolic functions, memory and learning), or months to years (e.g., affecting brain growth and brain plasticity, or emerging cognitive, behavioral, or social capacities)(*7*). Neurophysiological, neuroimaging, and observational studies have attempted to describe and quantify these early developmental changes, but there remains a need for non-invasive, objective biomarkers that can be measured serially across the months and years required for childhood development(*8-10*).

Human scalp hair, derived from the neuroectoderm and mesoderm, grows constantly at about 1 cm/month and evolves via prenatal lanugo, postnatal vellus, intermediate medullary, and terminal hair stages. Hair contains 65-85% proteins, 15-35% water, 1-9% lipids, and 0.1-5% pigments like melanin and trace elements(*11*). Constantly growing scalp hair incorporates both endogenous and exogenous proteins in a time-averaged chronological manner(*12*), unlike any other biospecimens(*13*). Therefore it is used for monitoring drug exposures, heavy metals, or other environmental toxins(*14*). Developmentally regulated hair proteins could offer biomarker candidates for brain development in early childhood.

However, all published data on hair proteins are limited to adult subjects, include relatively small sample sizes, and focus mainly on structural hair proteins. Lee et al. reported 343 hair proteins from three adults, showing evidence for post-translational modifications(*15*).

Laatsch et al. analyzed hair from 18 males and 3 females, reporting ethnic differences in keratins and keratin-associated proteins (KAPs)(*16*). Carlson et al. characterized hair proteins from one adult with limited sample availability(*17*), whereas Wu et al. used hierarchical protein clustering to match 10 monozygotic twin pairs and differentiate them from unrelated individuals(*18*).

Parker et al. reported quantifiable measures of identity discrimination and racial ancestry by detecting genetically variant peptides in the structural hair proteins for forensic purposes(*19*). To fill the extant gaps in knowledge, we analyzed non-structural hair proteins using ultra-performance liquid chromatography-tandem mass spectrometry (UPLC-MS/MS) in preschool children and their mothers. Our subjects were not exposed to early life adversity, as evidenced by parental income, household structure, health insurance, and parent education(*4*); all children were developmentally appropriate, healthy, and belonged to stable nuclear families (**Table S1**).

## RESULTS

### Features of hair proteins

Pooled hair samples from 40 children and 43 mothers were processed initially to identify non-structural hair shaft proteins, using protein purification to remove keratins. We subsequently analyzed individual hair samples from 8 mothers with 16 related children and 16 unrelated children, to create a master library of hair proteins with 3,124 proteoforms, representing the gene products of 2,278 genes. Expression of protein isoforms, alternative splicing of messenger RNA (mRNA), and post-translation modifications resulted in a higher number of hair proteins than their associated genes(*15, 20*). These data were deposited through the PRIDE repository(*21*) into the ProteomeXchange Consortium(*22, 23*).

Hair proteins observed in individual mothers and children contained 2,269 unique ‘proteoforms’ or protein isoforms; 1,438 proteins were commonly observed in mothers, 1,368 proteins were commonly observed in children, whereas 1,288 hair proteins showed individual variability among mothers and children. Higher spectral counts (p=0.0004) and higher numbers of proteins (p=0.001) were observed in mothers compared to children (**Fig.1**), perhaps reflecting a wider array of biological functions in adult females related to reproduction(*24-26*), aging(*27, 28*), or disease states(*29*). These age differences are explored further in subsequent analyses.

**Figure 1:**
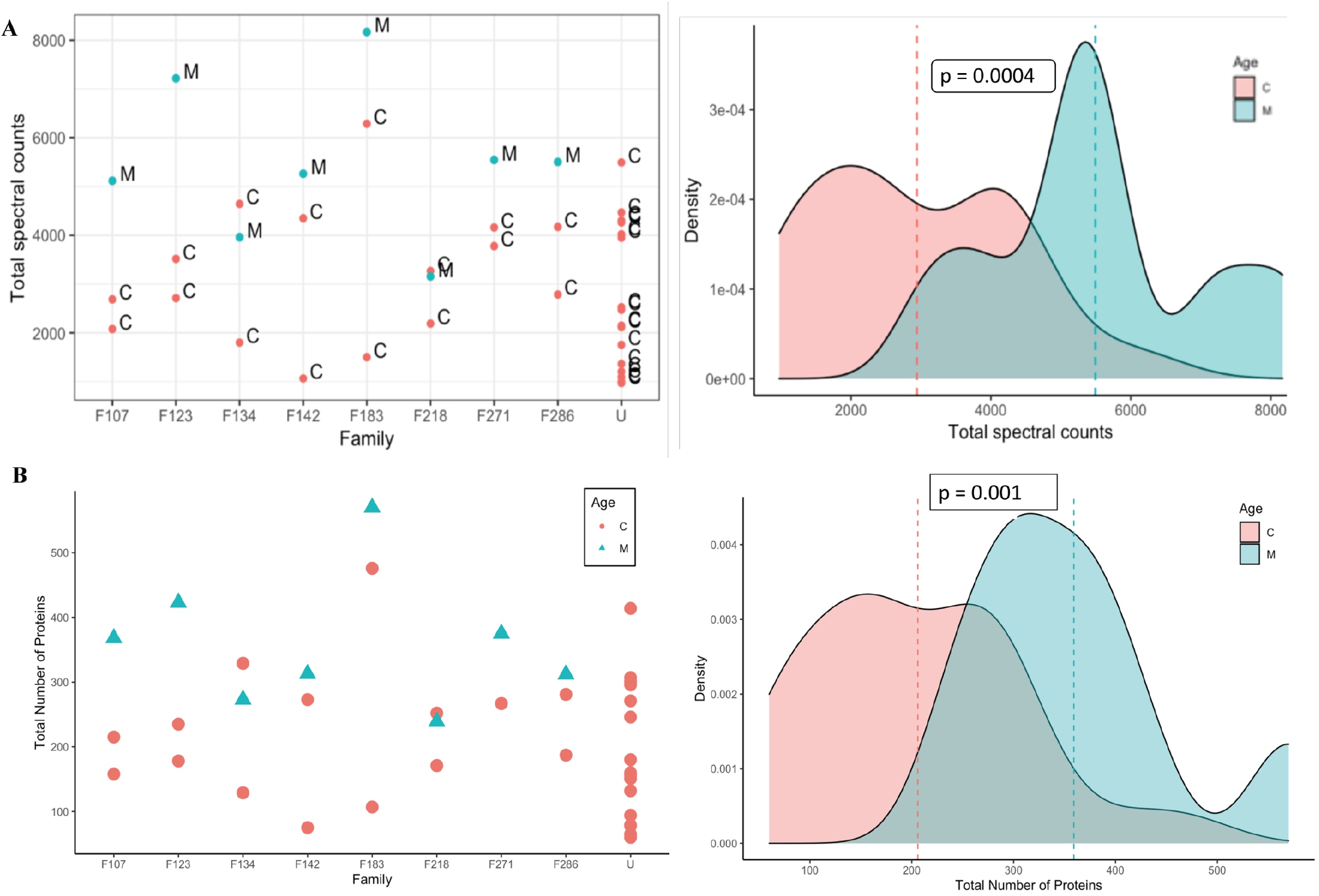
Hair proteins in mothers and children. **(A)** Protein spectral counts (p=0.0004) and **(B)** the numbers of proteins observed (with spectral counts >3) were consistently higher (p=0.001, Wilcoxon tests) in the mothers (M; Cyan) as compared to children (C; Pink). Mothers and their children (family labels: F107, F123, F134, F142, F183, F218, F271, F288) and Unrelated children (U) are identified on the X-axis: every mother except F134 and F218 had higher spectral counts and more hair proteins than her children.

The Uniprot database (https://www.uniprot.org/) revealed, for example, that 191 proteins observed in hair were regionally enriched in specific brain regions (based on the Allen Brain Atlas (https://human.brain-map.org/static/brainexplorer and the Human Brain Protein Atlas (https://www.proteinatlas.org/search/brain_category). Observed spectral counts for 2 brain proteins were higher in children (Lipocalin-1, Matrix metalloproteinase 7), 47 proteins were higher in mothers, 45 proteins lacked age-related differences, and 97 proteins were observed sporadically in individuals. Cellular functions and biological processes associated with the observed proteins were identified using PANTHER classification systems (**Fig.S1**, www.pantherdb.org).

### Hair protein profiles in individuals and families

Peptide spectral matches for each protein were combined to compare protein expression for all individuals and assess Spearman rank correlations. Hair proteins from the mothers were closely correlated with each other, whereas hair proteins in children showed correlations based on age and sex (**Fig.2A)**. Euclidean distances were calculated for pairwise comparisons between individuals (**Fig.2B**) and used for hierarchical clustering to identify subjects with similarities in the hair protein patterns (**Fig.2C**). Consistent with the correlation matrix, all mothers were clustered close together, younger children (0-2 years) were mostly located in one cluster, whereas older children were clustered with the mothers (**Fig.2C**). Boolean profiles of the hair proteins for each mother and her two children showed significantly shorter intra-family Manhattan distances (p<0.0002) as compared to 5000 ‘simulated’ families with mismatched mothers and children (**Fig.2D**), revealing a hereditary conservation of hair protein profiles in each family.

**Figure 2:**
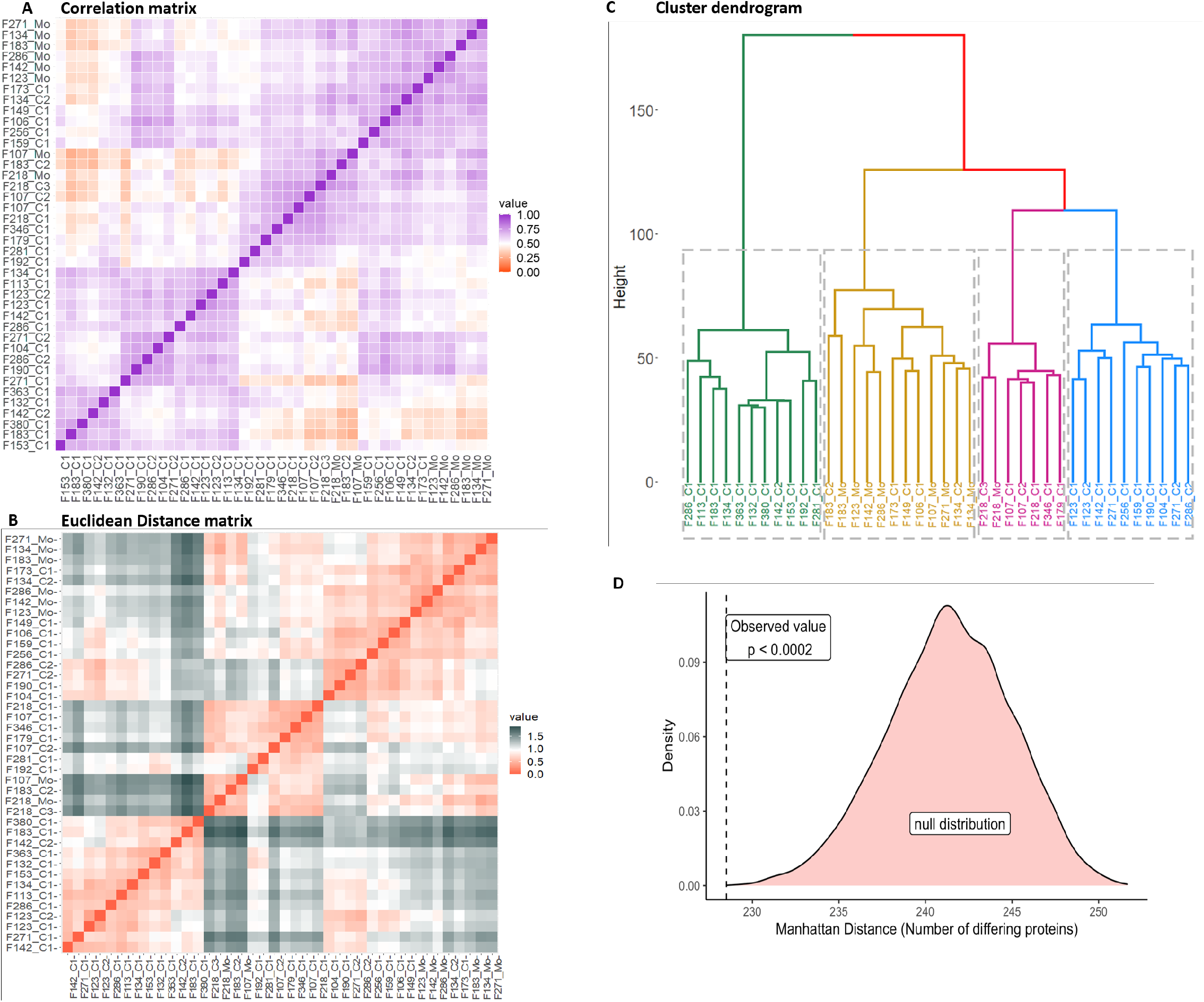
Similarities in hair protein profiles of individuals and families. **(A)** Spearman rank correlation matrix, with high (purple) to low (orange) correlation coefficients*; **(B)** Euclidean distances based on protein spectral counts, showing individuals more closely related (red) or more distant (grey) from each other*; **(C)** Hierarchical cluster dendrogram based on log spectral counts showing 7/8 mothers grouped in one cluster (mustard) with one mother in an adjacent cluster (pink); younger children (0-2 years) in one cluster (green) whereas older children dispersed in the other clusters*; **(D)** Intra-family Manhattan distances from Boolean hair protein profiles were shorter in mothers matched with their own children (p<0.0002) vs. 5000 simulated datasets created with mismatched mothers and children. *Note: Individuals are listed on the X- and Y-axes with their family identifier, with Mo for mother, C1 for younger child, C2 for older children in each family.

### Age- and sex-related differences in hair proteins

Principal Component Analysis (PCA)(*30-32*) and t-distributed Stochastic Neighbor Embedding (tSNE)(*33, 34*) were used to reduce the dimensionality of our data and to identify the major contributors of hair protein variability. Principal components 1-5 accounted for 61.6% of hair protein variability for all subjects, 57.5% for all children, 84.0% for all mothers, 60.8% for mothers and related children, and 62.3% for mothers and unrelated children.

***Age differences*** were observed by plotting the first two principal components (PC1, PC2) and tSNE dimensions (**Fig. 3**). We observed clusters of the younger children and mothers, with the older children dispersed across these groups (**Fig. 3A**). Similar clusters were observed from the remaining principal components. The tSNE projections also showed mothers located separately from the children (**Fig. 3B**). Proteins driving these differences showed higher spectral counts in mothers vs. children for SERPINB4 (serine protease inhibitor), POF1B (actin filament binder), PLEC (cytoskeleton binding protein), A2ML1 (α2-macroglobulin-like proteinase inhibitor), HIST1H3A (histone), UQCRQ (electron transfer from ubiquinol to cytochrome C), and AHCY (adenosylhomocysteine hydrolase). In contrast, mammaglobin-B (SCGB2A1), a heterodimerization protein that binds androgen and other steroids, was observed only in children (**Table S2**). Older children had higher spectral counts for PLEC (plectin), EIF3A (eukaryotic translation initiation factor 3), AHCY (adenosylhomocysteinase), HAL (histidine ammonia-lyase), and TUBA1C (Tubulin alpha 1c), whereas younger children had higher protein spectral counts for SCGB2A1 (secretoglobin 2A member 1) and CSN2 (casein beta) (**Table S3**).

**Figure 3:**
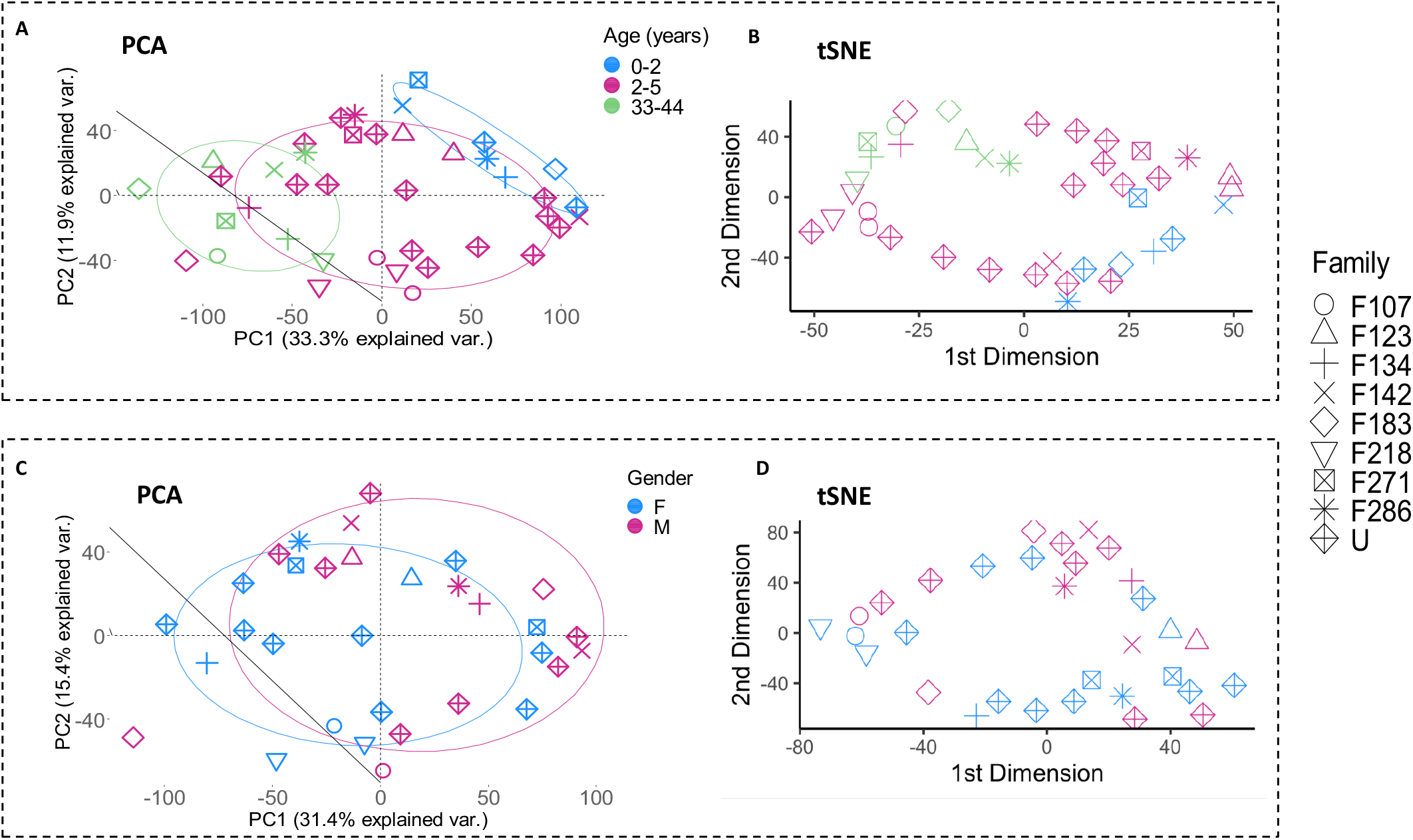
Age and sex-related differences in hair proteins. **(A)** First two principal components showing spatial separations by age, with children above 2 years (pink) located in between the children 0-2 years (blue, upper right) and the mothers (green, lower left). **(B)** The first two tSNE dimensions by age, showing mothers in the left upper quadrant separate from the children. Higher spectral counts for 7/17 hair proteins were observed in mothers (SERPINB4, POF1B, PLEC, A2ML1, HIST1H3A, UQCRQ, AHCY) and one protein (SCGB2A1) in children (Kruskal-Wallis and *post hoc* Benjamini-Hochberg corrections; see Table S2). **(C)** PCA analyses of all children showing overlapping circles for girls (blue) and boys (pink). **(D)** tSNE dimensions by sex, showing overlap between boys and girls. Higher spectral counts were observed only for CSN2 (Casein beta) in boys (p=0.0184) and ALMS1 (Alström syndrome protein 1) in girls (p=0.0214) (see Table S3).

***Sex differences*** showed slightly higher spectral counts in girls vs. boys (p=0.038) but no difference in the number of proteins (**Table S1**). PCA analyses and tSNE projections showed overlapping clusters of boys and girls (**Fig. 3C, 3D**). When comparing individual proteins, higher spectral counts were observed for CSN2 (Casein beta) in boys and ALMS1 (Alström syndrome protein 1) in girls (**Table S4**).

***Machine learning algorithms*** using Random Forest regressions(*35*) were used to predict age from their hair protein profiles. This model predicted age differences in mothers and children (R^2^=0.37, **Fig.4A**), but performance improved (R^2^=0.45) when predicting age for all children (**Fig.4B**). Random Forest classifier algorithm showed an acceptable mean accuracy for classifying mothers and children based on their predicted vs. observed age (mean area under the ROC curve = 0.93, **Fig.4C**; Wilcoxon test p=0.00011, **Fig.4D**).

**Figure 4:**
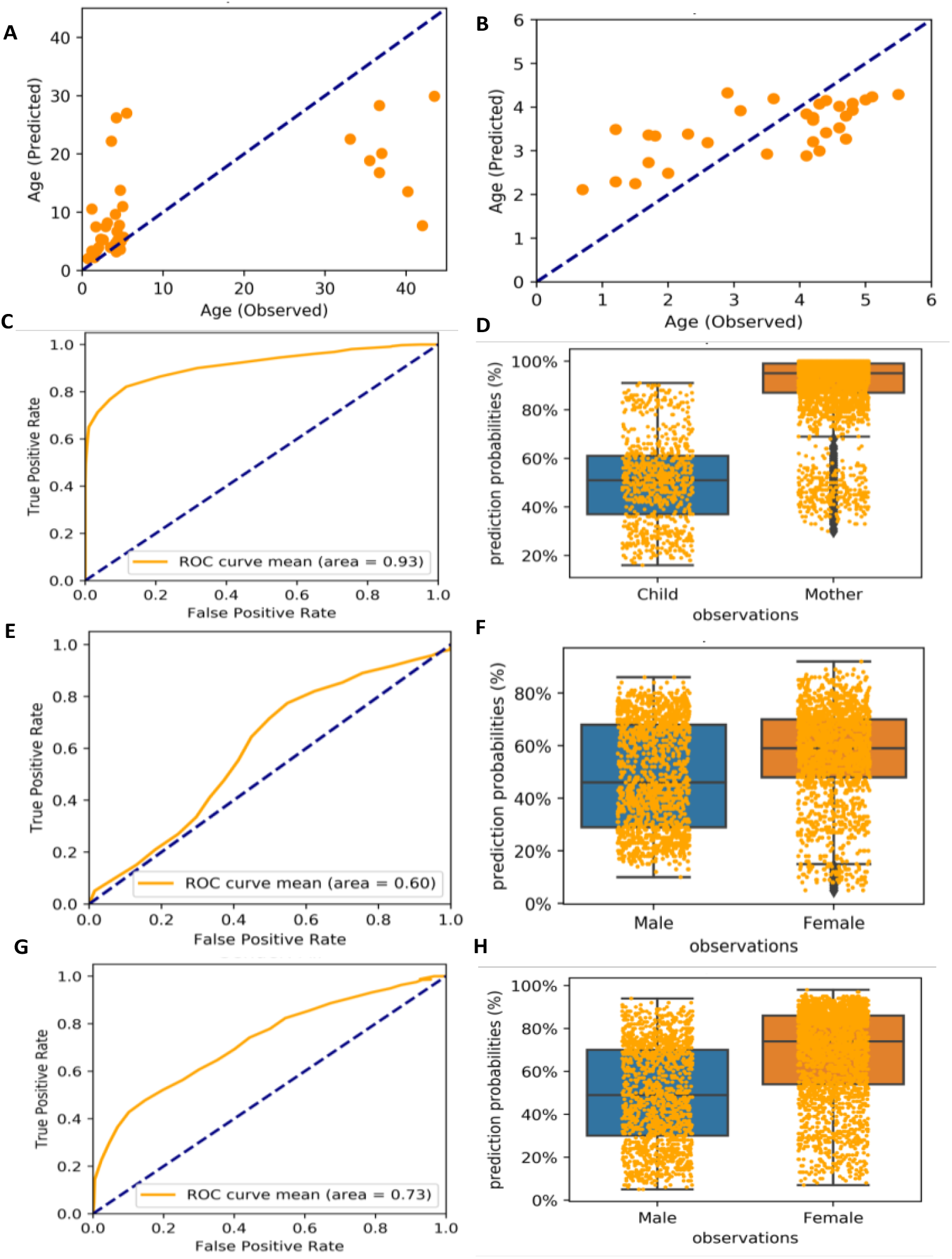
Machine learning algorithms predict age and sex from hair proteins. Mean scatterplot from 100 runs of Random Forest regression showing **(A)** observed vs. predicted age for mothers and children (R^2^ 0.37, p=0.00005) and **(B)** for all children (R^2^ 0.45, p=0.00004). **(C, D)** Random Forest plot showing mean accuracy for classifying mothers and children based on hair proteins (mean area under the ROC curve = 0.93, Wilcoxon test p=0.00011). **(E, F)** Random Forest plot showing mean accuracy for classifying by sex based on hair proteins for children (mean area under the ROC curve = 0.60, Wilcoxon test p=0.1703). **(G, H)** Random Forest plot improved when classifying all participants including mothers and children (area under the ROC curve = 0.73, Wilcoxon test p = 0.00831).

A Random Forest classifier to predict sex from hair protein profiles in children could not reliably differentiate boys from girls (mean area under the ROC curve = 0.6, **Fig.4E**; Wilcoxon test p = 0.1703; **Fig.4F**), but predictions improved when classifying all participants including mothers and children (area under the ROC curve = 0.73, **Fig.4G**; Wilcoxon test p = 0.0083, **Fig.4H**). The latter result is likely due to the age-based distinction between mothers and children, although sample size-related effects cannot be ruled out (25 vs. 17 females).

### The top contributors to hair protein variability

The top 5% proteins were identified as the most prominent contributors based on their total loading scores (TLS), explaining 64.3% of hair protein variability in all individuals, 89.5% in all mothers, 57.5% in all children, 49.3% in mothers and related children, and 64.6% in mothers and unrelated children (**Fig.5**). Keratins and KAPs are structural components of hair, but usually considered contaminants in most proteomics experiments, due to their high abundance in common lab contaminants. We therefore performed PCA analyses for all individuals with (**Fig.5A**) and without excluding the keratins and KAPs (**Fig.5B**). Structural proteins contributed to hair protein variability but with no differences in their biological significance. Separate PCA analyses performed to characterize the hair proteins observed in subpopulations of mothers (**Fig.5C**), children (**Fig.5D**), mothers and related children (**Fig.5E**), and mothers and unrelated children (**Fig.5F**) showed the same proteins ranked in all individuals and all children. Other than the histones, no other proteins were common between mothers and children. Tubulin alpha 1c (TUBA1C) was the only hair protein ranked within the top 5% in all groups, whereas PLEC, SERPINB4, and UQCRQ were observed in multiple groups.

**Figure 5:**
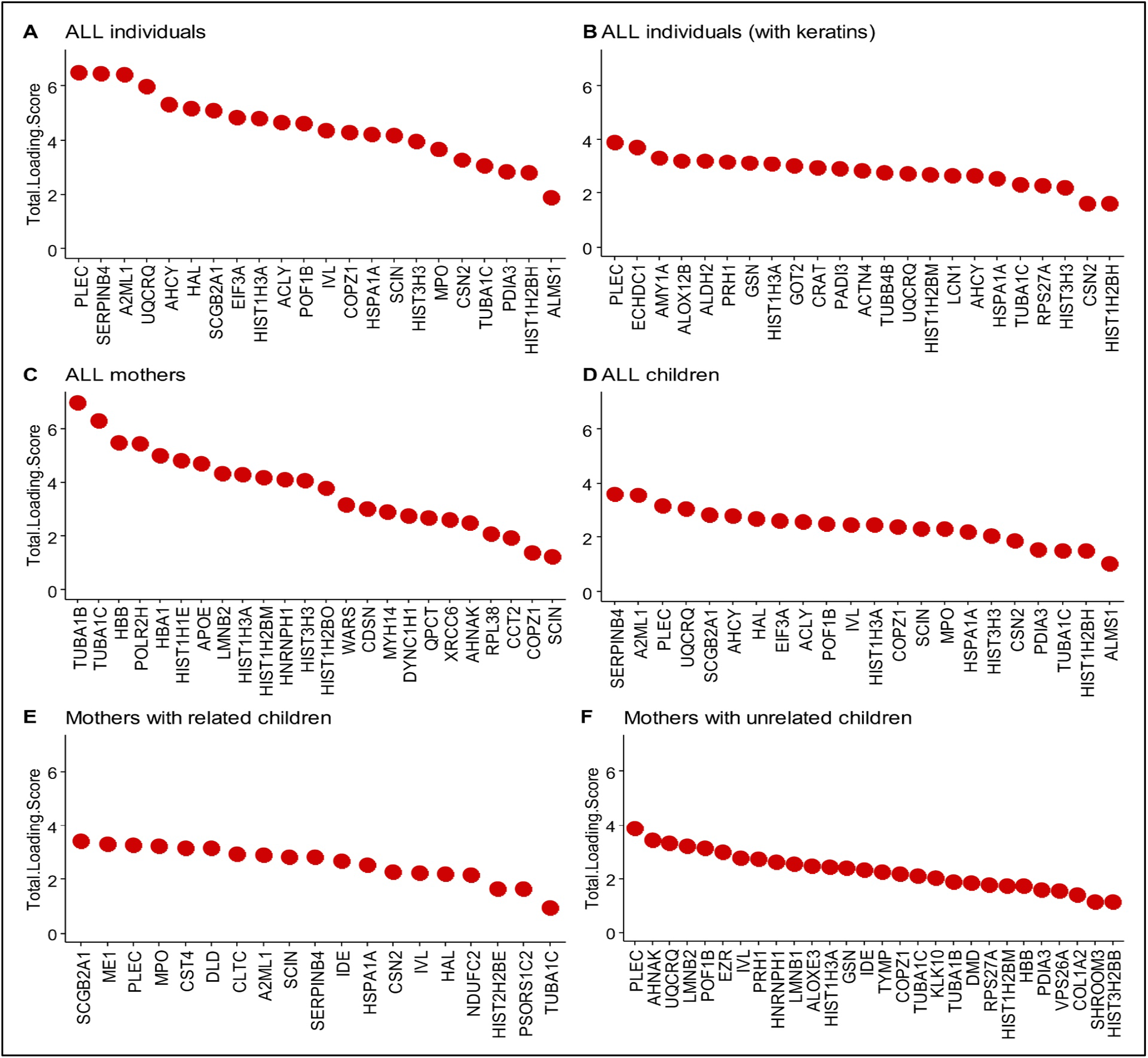
Top 5% proteins contributing to hair protein variability. The loading scores for each protein were weighted by the percent variance explained by each corresponding PC and then summed to give the Total Loading Score (TLS) for each protein. The top 5% proteins based on their TLS were identified as the most prominent contributors in each group. **(A)** All individuals (N=40, 49% of hair protein variability), **(B)** All individuals including keratins and KAPs (N=40, 64.3% variability); **(C)** All mothers (n=8, 89.5% variability); **(D)** All children (n=32, 57.5% variability); **(E)** Mothers (n=8) and their biological children (n=16) (49.3% variability), and **(F)** Mothers (n=8) and unrelated children (n=16) (64.6% variability).

### Biological role(s) of the strongest contributors to hair protein variability

Based on experimentally observed human data in the Ingenuity Knowledge Base, log-fold-change values of the top 5% proteins from our dataset were used to analyze direct and indirect relationships between molecules. Protein networks for the top 5% hair proteins contributing to age-related differences between mothers and children (**Fig.6**) and similar analyses for sex-related differences between girls and boys were examined (**Fig.7**). Using these molecular relationships as input for Ingenuity Pathway Analysis, we found proteins involved in *cellular metabolism* including the protein ubiquitination pathway, Sirtuin signaling pathway, 14-3-3 mediated signaling, Wnt-Ca^++^ pathway, histidine degradation, mitochondrial function, and oxidative phosphorylation (**Fig.8**). Other proteins were associated with *immune responses*, including phagosome maturation, IL-8 signaling, and regulation of macrophages, fibroblasts, and endothelial cells, or involved in the regulation of *stress-related pathways*, including corticotropin releasing hormone signaling, glucocorticoid receptor signaling, prolactin and aldosterone signaling. Finally, hair proteins associated with *brain development* including axonal guidance and gap junction signaling were also identified (**Fig.8**).

**Figure 6:**
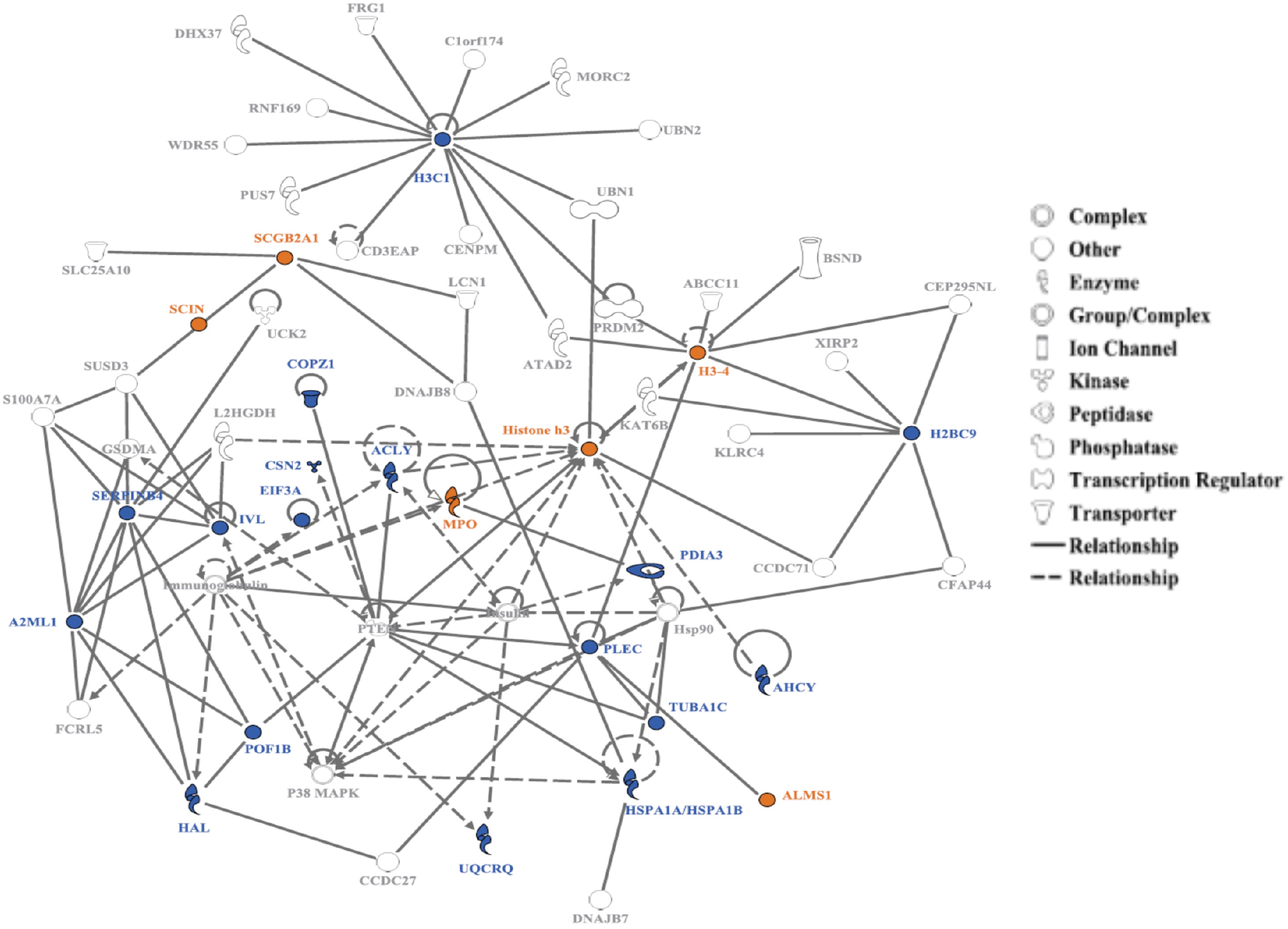
Protein network for the top 5% hair proteins contributing to age-related differences between mothers and children. Some hair proteins had higher spectral counts in children (Orange) and others had higher spectral counts in mothers (Blue); continuous and interrupted lines show direct and indirect relationships, respectively. Mothers show higher protein spectral counts mostly for the ‘enzymes’ and ‘peptidases’ involved in cellular and metabolic processes, while proteins with higher spectral counts in children belong to ‘other’ group involved in growth and biological regulation.

**Figure 7:**
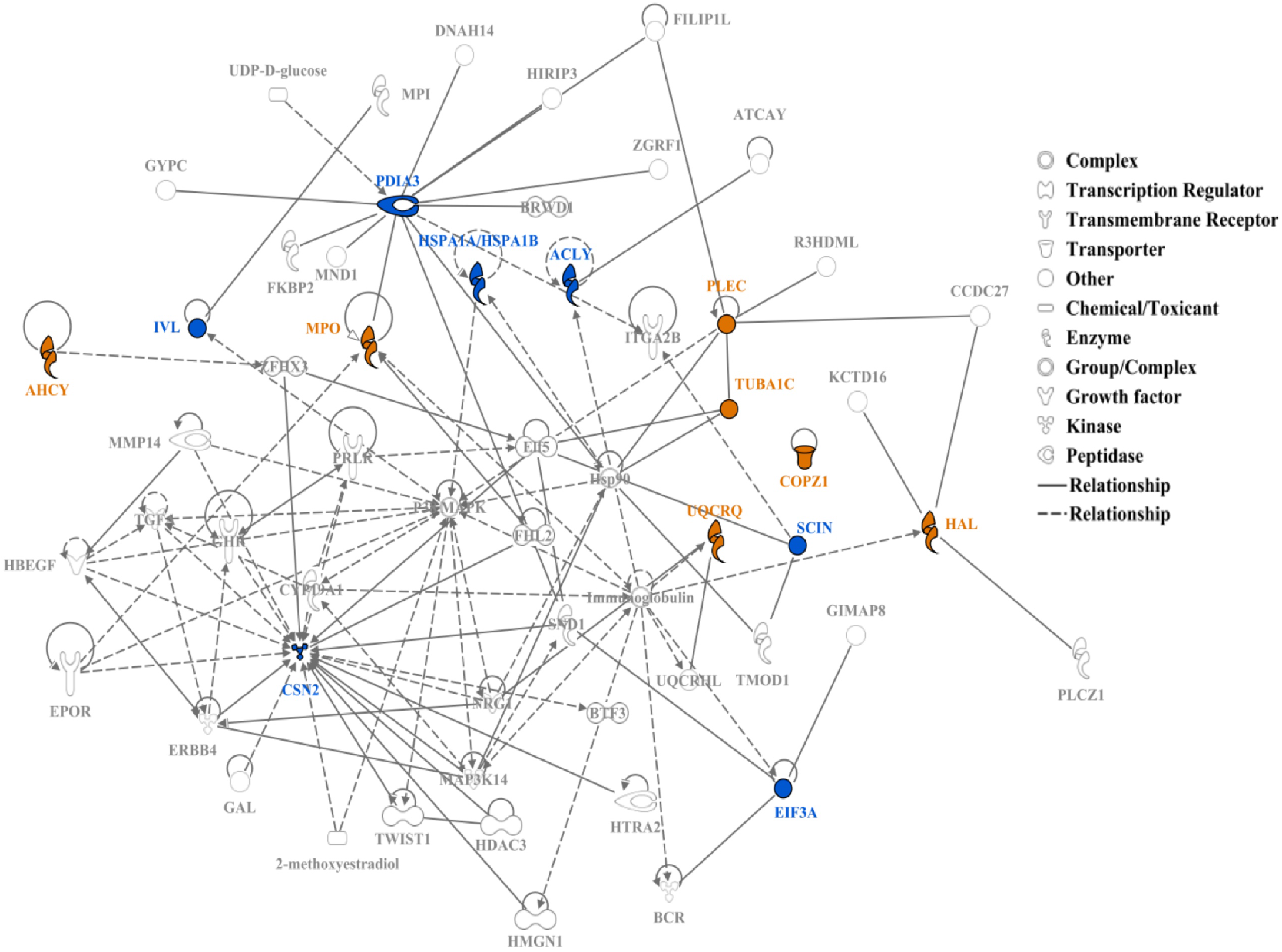
Protein network for top 5% hair proteins contributing to sex differences between boys and girls. Some proteins had higher spectral counts in girls (Orange) and others had higher spectral counts in boys (Blue); continuous lines show direct relationships and interrupted lines denote indirect relationships. Girls show higher protein spectral counts mostly for ‘enzymes’ or ‘transporters’ associated with cellular localization and metabolic processes. Proteins with higher spectral count in boys are ‘enzymes’ like ‘kinases’ or ‘peptidases’ associated with cellular and metabolic processes and biological regulation.

**Figure 8:**
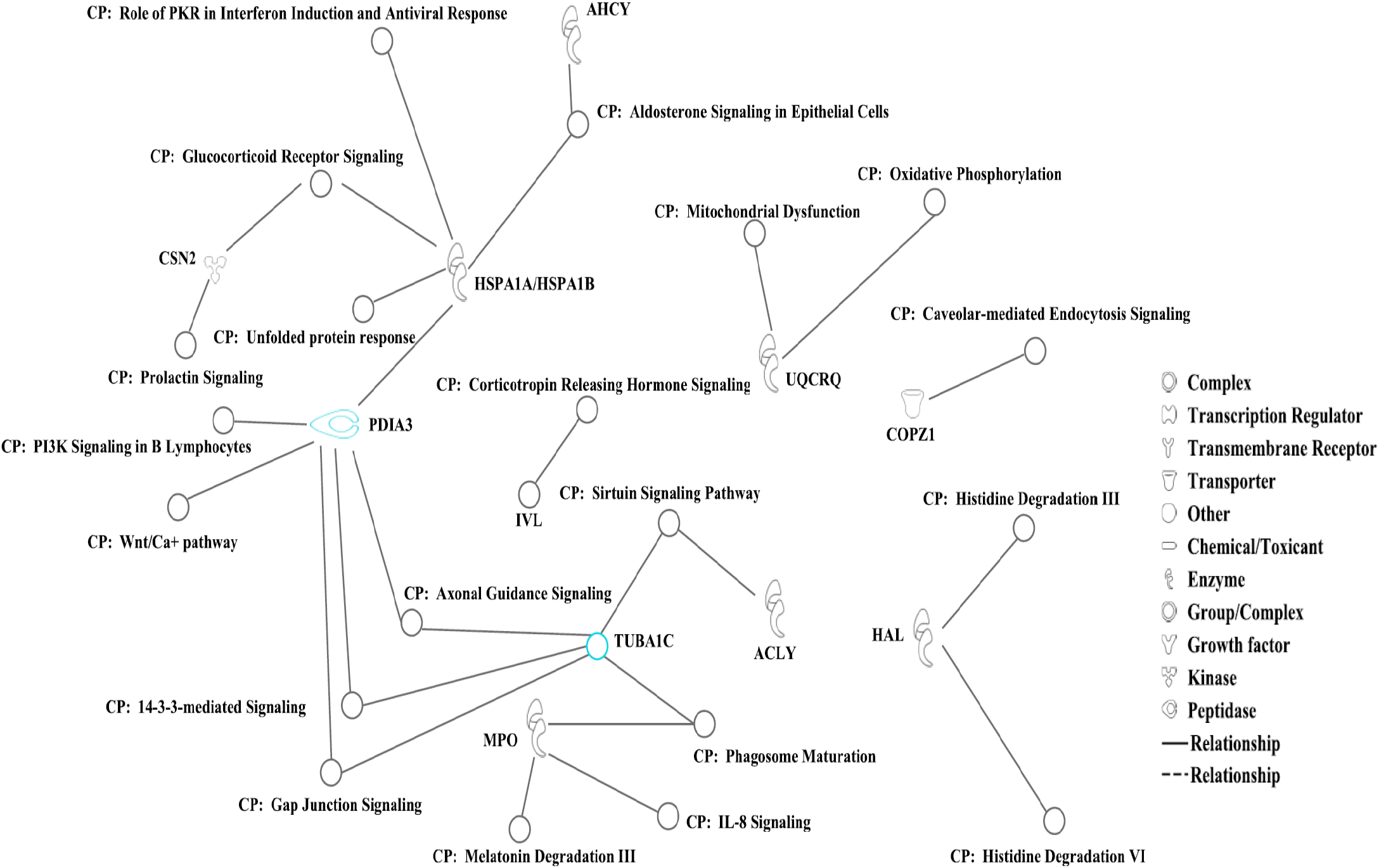
Canonical pathways associated with biologically significant proteins from the top 5% variables in all individuals (n = 40) contributing to age- and sex-related differences were identified with Ingenuity Pathway Analysis. Most of these proteins are involved in cellular metabolism, immune responses, brain development, and regulation of stress-related pathways.

## DISCUSSION

Growing hair provides a time-averaged molecular record unavailable from any other tissue or biospecimens(*13*). The hair follicle includes most cellular processes, including cell growth, cell death, interactions of histologically different cell types, cell migration and differentiation, and it responds to multiple hormones, microenvironmental and systemic changes(*36*). Structural and functional characteristics of hair follicles also mimic the renal tubules, potentially performing unique, distributed excretory functions(*37*).

The chemical composition of hair(*11, 38-42*) and its structural proteins (keratins, KAPs) are well-studied(*16-19*), but with minimal data on non-structural hair proteins. We found 2,269 non-structural hair proteins with important differences between mothers and children, age- and sex-related differences among preschool children, and conservation of protein profiles within families. Hair proteins driving variability in different populations were found to play vital roles in functions other than those of trichocytes in the hair follicle, including cellular metabolic pathways, brain development, immune signaling, and stress regulation. Thus, unbiased or targeted protein profiles from single or serial hair samples (or sequential hair segments) could be used as probes for child development(*43, 44*) or life-course studies(*29, 45, 46*).

We observed age-related hair protein profiles in children and mothers, with distinct patterns emerging in multiple analyses. Differences between mothers and children were largely driven by increased maternal expression of SERPINB4, PLEC, and UQCRQ. SERPINB4 is a granzyme inhibitor linked with squamous cell carcinomas and chronic liver disease(*47-49*), Plectin mutations were linked with epidermolysis bullosa simplex, but has also emerged as novel susceptibility gene for testicular germ cell tumors(*50-52*), and UQCRQ is a nuclear protein in the mitochondrial respiratory chain complex III essential for normal brain development(*53*).

Mammaglobin-B (SCGB2A1), which is linked with familial febrile seizures in preschool children(*54, 55*) and chemoresistant cancers in adults(*56*), was observed only in children’s hair.

Our analyses showed minimal sex differences in early childhood, confirmed by Random Forest predictive models. Biological pathways for cellular metabolism and innate immunity appeared more prominent in girls, whereas brain development and regulation of stress responses appeared more prominent in boys. Perhaps sex differences in hair proteins may be accentuated following the onset of puberty(*57*). Although hair protein profiles were conserved in mothers and their biological children, future studies in mother-child dyads and monozygotic vs. dizygotic twins are required to quantify the heritability of hair proteins(*58*).

Experimental human data from the Ingenuity Pathway Analysis identified hair proteins associated with axon guidance(*59*) and gap junction signaling(*60*). For example, Tubulin alpha 1C was identified from all sub-populations in our dataset. TUBA1C (on chromosome 12) is amongst the most over-expressed brain-specific genes, forming a major component of the microtubules that determine neuronal shape and function, and is typically used as a marker of neural differentiation(*61*). We observed 191 hair proteins regionally enriched in the brain and speculate that hair proteomics could complement neuroimaging and neurophysiological data on early brain development(*8*). For example, specific plasma proteins were associated with higher non-verbal intelligence and pro-inflammatory proteins associated with lower intelligence in children from Nepal(*62*). That study used an FDR of 5%, whereas our FDR threshold was set at 1% or lower.

Future developmental studies with larger sample sizes could correlate hair proteins with cognitive or behavioral outcomes, thus investigating their role in normal and altered brain development(*63*). Clinical trials of novel therapeutics to enhance brain recovery, in stroke or traumatic brain injury for example, would also benefit from brain-region specific biomarkers. *“Despite intensive search, there is no such biomarker in brain imaging or serum”(64)* and therefore, clinical trials of neural repair therapeutics have suffered from reliance on insensitive behavioral outcomes(*64*).

These findings must be interpreted in the light of three limitations. First, our sample size of 32 children was insufficient to examine developmental differences at each age in the preschool period. We selected healthy children from homogenous socioeconomic environments; they did not experience any adverse conditions and therefore, our data may not represent the full range of hair protein profiles present in the general population. Our sample size, however, is larger than most other studies of hair proteomics in adults and it is the first to include mothers and children. Our study design also allowed us to investigate differences in hair protein profiles between related and unrelated individuals, as well as differences between adults and children.

Second, our proteomics platform relied on peptide spectral matches, which presented only semi-quantitative data on the abundance of hair proteins between individuals. Since this was the first-ever study investigating non-structural proteins from hair in humans, we chose a ‘shot-gun’ proteomics approach rather than targeted and more quantitative approaches. Now that we have established large libraries of hair proteins for mothers and children, future studies can be designed for the quantitation of specific protein targets or protein groups. Lastly, we did not correlate hair proteins with the child’s developmental milestones or their cognitive and behavioral data. We feel that the sample size limitations at each age would preclude any generalizable conclusions from such analyses.

Despite these limitations, our initial findings reveal the potential role of non-structural hair proteins as biomarkers for brain development, metabolic, immune, or stress-regulatory pathways, providing a rich source of chronologically-ordered information for life-course studies and early childhood development.

## MATERIALS AND METHODS

After IRB approval and parental consent, mothers and children aged 1-6 years were enrolled from local preschool facilities. We excluded children with tinea capitis, alopecia areata, eczema, or other scalp conditions; those receiving any prescription or over-the-counter drugs; or steroid therapy in the past 3 months; or those with chronic medical conditions, developmental delay, or chemical exposures to hair prior to study entry. Hair samples from the posterior vertex area were trimmed at 0.1mm from the scalp, weighed, and stored in Ziploc® bags at 4°C.

### Hair extraction

Finely chopped hair (10 mg) was incubated in 500μl Tri-Reagent® (Molecular Research Center, Inc; Fisher Scientific) at room temperature (RT) for 10min, followed by 100μl chloroform (shaken vigorously for 15sec, stored for 10min). Samples were centrifuged at 12000g, 4°C for 10 min to separate the aqueous RNA fraction. Samples were incubated with 1.5ml acetone at –30°C for 1 hour, followed by centrifugation at 8000g for 5 min to sediment protein. The supernatant was removed, the sediment suspended in 1ml, 0.3 M guanidine hydrochloride/95% ethanol/2.5% glycerol, stored for 10min at RT, followed by centrifugation at 8000g for 5 min. The supernatant was decanted and the wash repeated twice. The protein pellet was washed twice in 95% ethanol (without glycerol), centrifuged for 5min, 8000g at 4°C and the pellet stored frozen under fresh 95% ethanol at –20°C.

### Proteomics methods

Samples were brought to RT, centrifuged at 8000g for 5 min, the ethanol decanted and dried under vacuum. Protein pellets were resuspended in 50mM ammonium bicarbonate in the presence of 0.0015% ProteaseMAX (Promega) and total protein amount was estimated with Pierce BCA assays (Thermo Fisher Scientific). Proteins were digested with 0.25µg of Trypsin/LysC (Promega) at a 1:100 enzyme/substrate ratio overnight at 37°C. Proteolytic digestion was quenched with 1% formic acid; peptides were dried by speed vac before dissolving in 30µl of reconstitution buffer (2% acetonitrile + 0.1% Formic acid) to a concentration of 1 µg/µl; 2µl of this solution was injected into the MS instrument.

Experiments were performed on the Orbitrap Fusion Tribrid mass spectrometer (Thermo Scientific) coupled with ACQUITY M-Class ultra-performance liquid chromatography (UPLC, Waters Corporation). For a typical LCMS experiment (Liquid Chromatography/Mass Spectrometry), a flow rate of 450 nL/min was used, where mobile phase A is 0.2% formic acid in water and mobile phase B is 0.2% formic acid in acetonitrile. Analytical columns were pulled using fused silica (I.D. 100 microns) and packed with Magic 1.8-micron 120Å UChrom C18 stationary phase (nanoLCMS Solutions) to a length of ∼25 cm. Peptides were directly injected onto the analytical column using a gradient (2-45% B, followed by a high-B wash) of 80 minutes. The MS was operated in data-dependent fashion using CID (collision induced dissociation) for generating MS/MS spectra, collected in the ion trap with collisional energy set at 35.

The *.RAW data files were processed using Byonic v3.2.0 (ProteinMetrics) to infer protein isoforms using the Uniprot *homo sapiens* database. Proteolysis with Trypsin/LysC was assumed to be semi-specific allowing for N-ragged cleavage with up to 2 missed cleavage sites. Precursor mass accuracies were held within 12 ppm and 0.4 Da for MS/MS fragments. Proteins were held to a false discovery rate (FDR) of 1% or lower, using standard target-decoy approaches(*65*), and only the proteins with >3 spectral counts were selected for further data processing; keratins and KAPs were removed at this stage.

### Statistical analysis

Spectral counts were used to calculate Euclidean distances between individuals; Euclidean distances were used for hierarchical clustering. A correlation matrix with Spearman’s coefficient was also used for rank-based depiction of similarities between the individual hair proteomes.

Principal Component Analysis (PCA) was used to reduce dimensionality of the dataset. PCA is a widely used technique for data analysis and modeling(*30*) of linear combinations of the original dimensions called principal components(*32*). The largest proportion of data variance is captured by the first principal component, the second largest proportion of variance falls along the second principal component, and so on(*31*). For the first five principal components from each PCA, we multiplied the loading scores for each protein by the percent variance explained by that corresponding principal component; the weighted scores were summed for each protein to give its Total Loading Score (TLS).

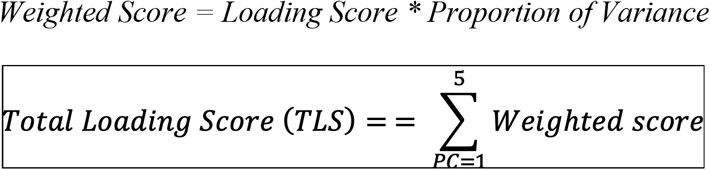

Based on their TLS values, top 5% proteins were selected as the main drivers of variability in hair protein expression.

Additionally, we used t-distributed stochastic neighboring embedding (tSNE), a non-linear probabilistic approach(*33, 34*), to visualize proteins with non-linear similarity in high-dimensional space as neighbors in low-dimensional linear depictions. Unlike the reproducible PCA results, the probabilistic nature of tSNE can result in somewhat different results with each computation. To avoid serendipitous results, we ran each computation at least 10 times to ensure reproducibility. For each computation, the maximum number of iterations to converge was set to 1000, and perplexity set to the maximum permitted value. Statistical significance of tSNE clusterings was calculated by how often a given statistic was reproduced in 1000 simulations of permuted versions of the dataset.

Boolean profiles of the hair proteins were also compared between the original dataset (each mother with her own children) and 5000 simulated datasets, created by swapping mothers between families such that no mother was paired with her own children, but the two siblings remained together in all simulated datasets. Observed conservation in pairwise intra-family Manhattan distances from the original dataset could then be attributed to the similarities in hair protein expression between each mother and her children.

For the top 5% proteins in children (n=32), we averaged spectral counts for girls and boys separately, and divided the girls’ average by the boys’ average. Resulting values were converted to log-base 2. The same process was followed for spectral counts from mothers and children.

Log fold-change values of the top 5% proteins were used as input for Ingenuity Pathway Analysis (Qiagen: https://digitalinsights.qiagen.com/products/features/). We analyzed direct and indirect relationships between molecules based on experimentally observed data, restricted to human databases in the Ingenuity Knowledge Base.

We used Random Forest (RF) models for both the classification (boy vs. girl, mother vs. child) and regression (age prediction) tasks, with protein concentrations as model features and individuals as samples(*35*). In classification, the model output was the probability of an individual being female (sex classification), or being a mother (person classification). For regression (age prediction), the model output was the individual’s predicted age.

Results were based on a 10-fold cross-validation repeated 100 times. Members of the same family were included in the same set, i.e. either training or test sets, to avoid information leak due to familial similarities. For the age prediction, we evaluated results using the R^2^ coefficient of determination and the linear model p-value fitted on the predicted and observed data. For the classification tasks, we used area under the ROC curve (AUC) and the Wilcoxon-Mann-Whitney-test, testing the null-hypothesis that one distribution is not stochastically greater than the other.

## Acknowledgements

The authors sincerely acknowledge the leadership of Santa Clara County Office of Education, the children, parents and staff at Bing Nursery School, Stanford Arboretum Children’s Center, Children’s Center of the Stanford Community Stanford Madera Grove Children’s Center, McKinley, Rouleau, and Wool Creek Head Start Centers for their partnership. We also thank Drs. David K. Stevenson, Gary Shaw, and Timothy Cornell for their valuable suggestions on previous versions of this manuscript; Sahil Tembulkar, Jitka Hiscox, and Marcela Lopez for help with collecting or processing the hair samples, Tanida Plamintr McHatton for administrative assistance, and Grant Padia for financial management.

## Funding

The Maternal & Child Health Research Institute (MCHRI) at Stanford, the *Eunice Kennedy Shriver* National Institute for Child Health & Human Development (R01 HD099296), the National Cancer Institute (P30 CA124435) for the Stanford Cancer Institute Proteomics/Mass Spectrometry Shared Resource, and the National Institute of General Medical Sciences (R35 GM138353) supported this research. Study sponsors had no role in the design or conduct of the study; the collection, management, analysis, or interpretation of the data; the preparation, review, approval, or decision to publish this manuscript.

## Author Contributions

KJSA designed the study and obtained grant funding, GT obtained hair samples, CRR performed the protein extractions, KS and RDL performed the proteomics experiments, KS, RDL, CRR and KJSA wrote initial drafts and edited the manuscript; KS, RDL, DD, MX and NA performed statistical analyses and created the figures; all authors reviewed and made critical revisions of the manuscript and approved the final version to be published.

## Competing interests

The authors received no honoraria, grants, or other payments for writing this manuscript and they report no relevant financial relationships and no conflicts of interest.

## SUPPLEMENTARY MATERIALS

**Table S1:** Demographic Characteristics and Hair Protein Data for Mothers and Children

| Family Code | Subject | Age (months) | Age (years) | Gender | Race | Ethnicity | # of hair proteins | Peptide Spectral Matches (PSMs) |
| --- | --- | --- | --- | --- | --- | --- | --- | --- |
| **F107** | Mother | 450.6 | 37.6 | F | White | Non-Hispanic | 819 | 6533 |
|  | Child1 | 27.6 | 2.3 | F | White | Non-Hispanic | 568 | 3949 |
|  | Child2 | 58.0 | 4.8 | M | White | Other | 464 | 2873 |
| **F123** | Mother | 447.1 | 37.3 | F | White | Non-Hispanic | 809 | 10370 |
|  | Child1 | 24.0 | 2 | F | White | Non-Hispanic | 499 | 3728 |
|  | Child2 | 52.4 | 4.4 | M | White | Non-Hispanic | 573 | 5078 |
| **F134** | Mother | 431.8 | 35.9 | F | White | Missing | 684 | 5445 |
|  | Child1 | 20.9 | 1.74 | M | Mixed | Missing | 387 | 2760 |
|  | Child2 | 67.6 | 5.6 | F | Mixed | Missing | 759 | 6872 |
| **F142** | Mother | 447.3 | 37.3 | F | Asian | Missing | 650 | 8370 |
|  | Child1 | 20.1 | 1.7 | M | Mixed | Missing | 581 | 6208 |
|  | Child2 | 50.6 | 4.2 | M | Mixed | Missing | 226 | 2353 |
| **F183** | Mother | 530.0 | 44.2 | F | Asian | Non-Hispanic | 1090 | 10527 |
|  | Child1 | 8.5 | 0.7 | M | Asian | Non-Hispanic | 314 | 2331 |
|  | Child2 | 44.0 | 3.7 | M | Asian | Non-Hispanic | 1010 | 8065 |
| **F218** | Mother | 504 | 42 | F | White | Hispanic | 609 | 4144 |
|  | Child1 | 35.2 | 2.9 | F | White | Hispanic | 631 | 4475 |
|  | Child2 | 58.5 | 4.9 | F | White | Hispanic | 524 | 3107 |
| **F271** | Mother | 402.6 | 33.6 | F | White | Non-Hispanic | 769 | 7525 |
|  | Child1 | 15.1 | 1.3 | F | White | Non-Hispanic | 557 | 7161 |
|  | Child2 | 42.5 | 3.5 | F | White | Non-Hispanic | 600 | 5615 |
| **F286** | Mother | 489.8 | 40.8 | F | White | Non-Hispanic | 616 | 9209 |
|  | Child1 | 22.0 | 1.8 | M | White | Non-Hispanic | 403 | 4727 |
|  | Child2 | 52.5 | 4.4 | F | White | Non-Hispanic | 614 | 6061 |
| **F346** | Child | 50.3 | 4.2 | M | White | Missing | 475 | 3429 |
| **F192** | Child | 38.1 | 3.2 | F | Other | Hispanic | 283 | 1731 |
| **F132** | Child | 51.4 | 4.3 | F | White | Missing | 272 | 1892 |
| **F363** | Child | 56.6 | 4.7 | M | Mixed | Non-Hispanic | 270 | 1914 |
| **F281** | Child | 53.5 | 4.5 | M | Mixed | Mixed | 406 | 3192 |
| **F173** | Child | 51.3 | 4.3 | F | Other | Other | 835 | 7168 |
| **F380** | Child | 14.8 | 1.2 | M | Asian | Missing | 237 | 1814 |
| **F159** | Child | 62.7 | 5.2 | F | White | Non-Hispanic | 485 | 3830 |
| **F179** | Child | 53.0 | 4.4 | F | Asian | Other | 494 | 2926 |
| **F149** | Child | 61.3 | 5.1 | F | Mixed | Non-Hispanic | 698 | 5733 |
| **F106** | Child | 56.7 | 4.7 | F | Asian | Non-Hispanic | 668 | 6390 |
| **F153** | Child | 57.1 | 4.8 | M | Asian | Missing | 275 | 2549 |
| **F256** | Child | 55.8 | 4.7 | M | Mixed | Mixed | 638 | 7016 |
| **F190** | Child | 31.5 | 2.6 | F | Asian | Other | 527 | 7460 |
| **F104** | Child | 50.2 | 4.2 | M | White | Non-Hispanic | 672 | 8084 |
| **F113** | Child | 17.9 | 1.5 | F | White | Non-Hispanic | 441 | 3640 |

Demographic data, total number of proteins, and peptide spectral matches observed in all 40 individuals. Mothers' hair (n=8) had significantly higher number of proteins ( $p=0.001$ ) and protein spectral matches ( $p=0.0004$ ) compared to children's hair (n=32). Related children (n=16, Cyan) are grouped with their mothers and unrelated children are listed below (n=16, Orange).

**Table S2:**
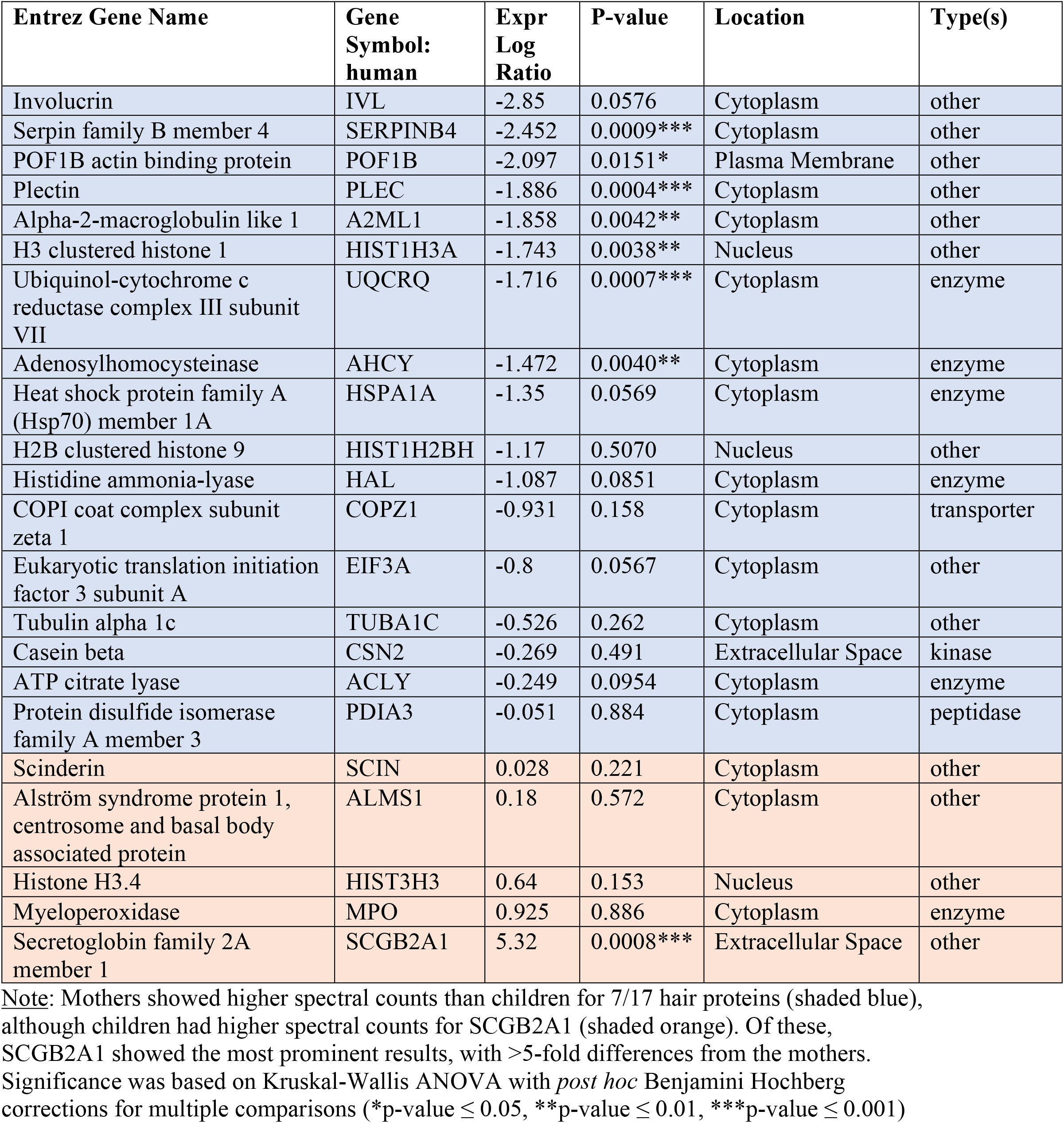
Hair Proteins mediating differences between Mothers and Children

**Table S3:**
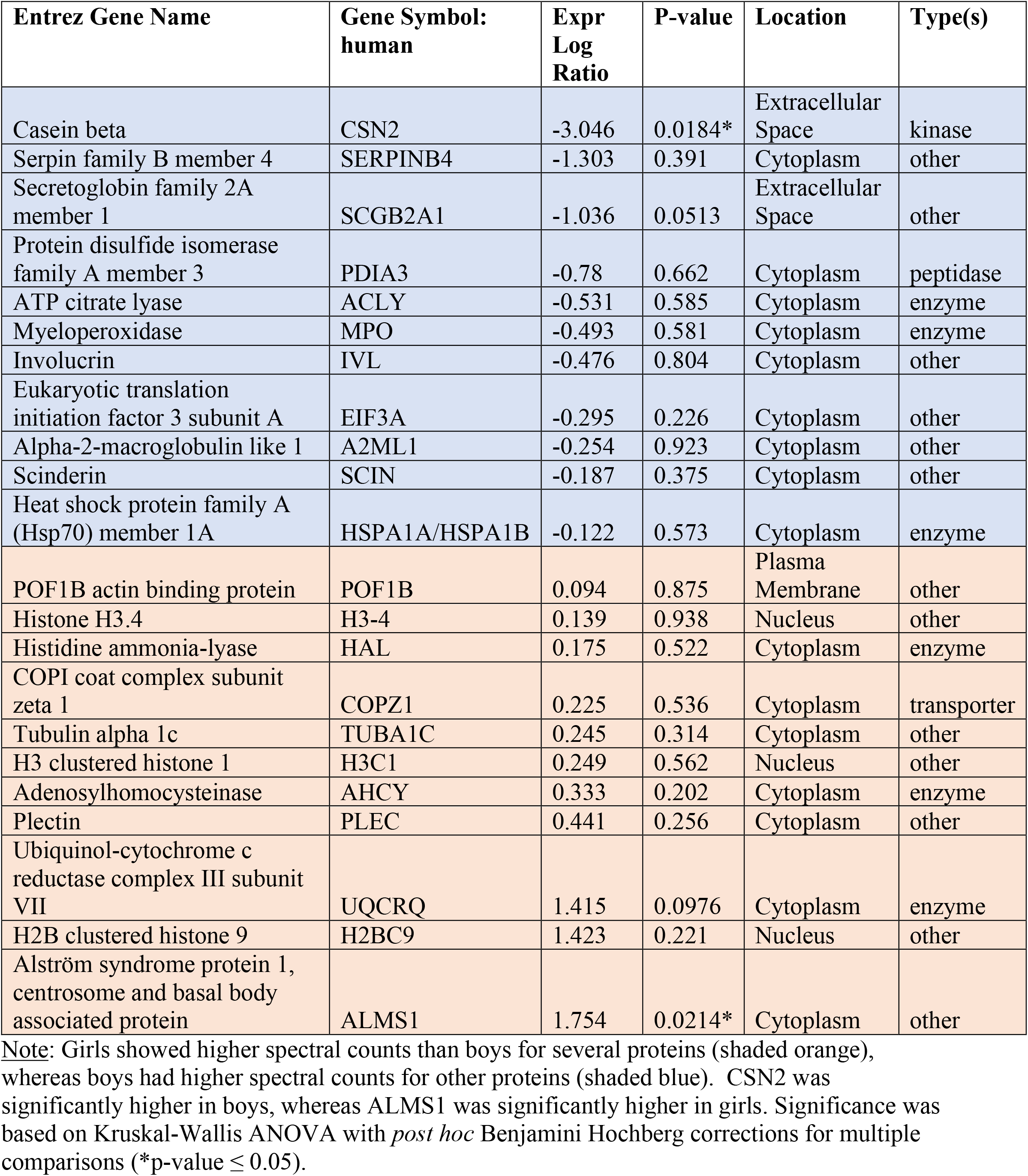
Hair proteins mediating differences between Boys and Girls

**Figure S1:**
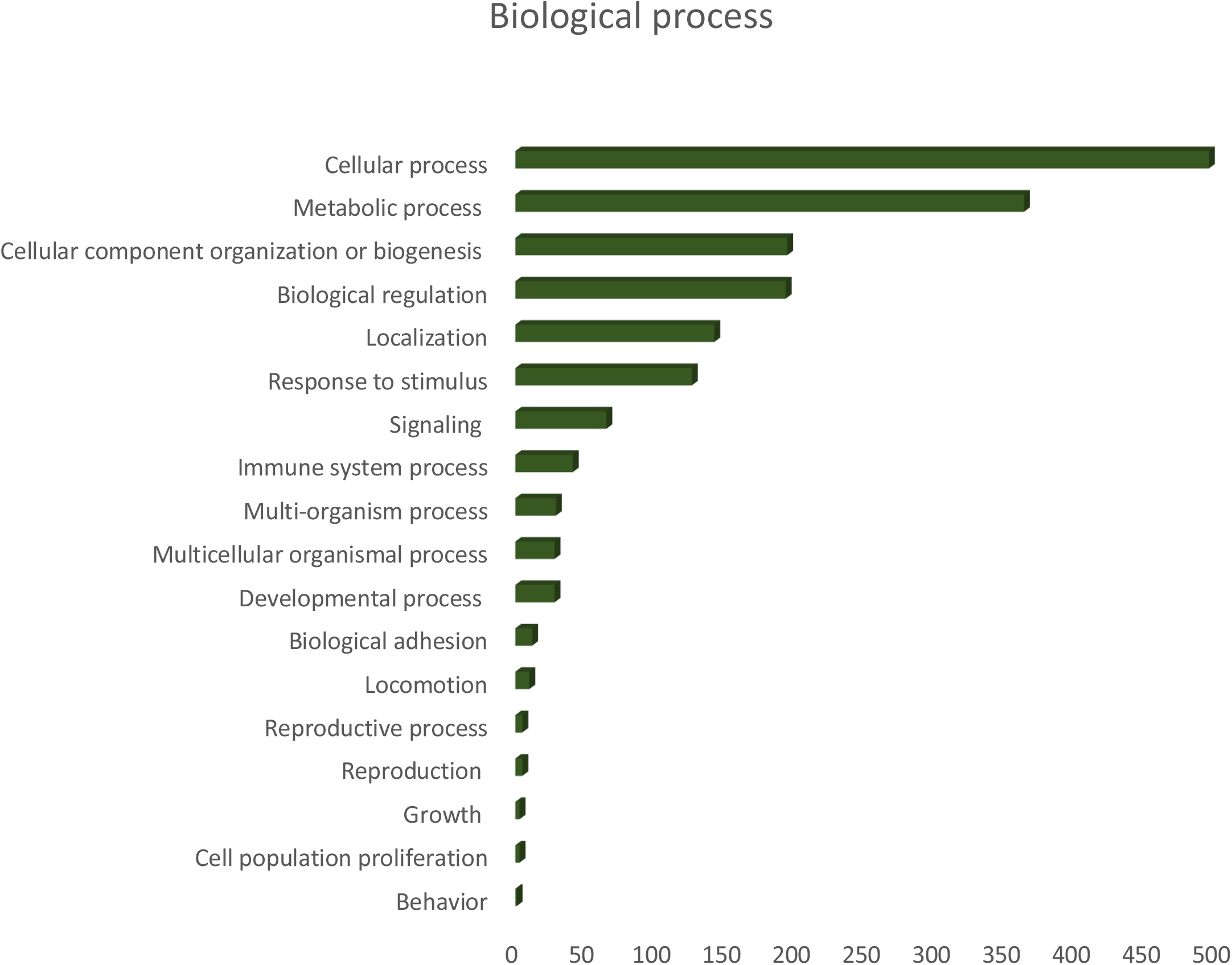
Number of proteins associated with specific biological processes identified by PANTHER classification system. Most of the proteins are associated with cellular functions and metabolic processes.

**Figure S2:**
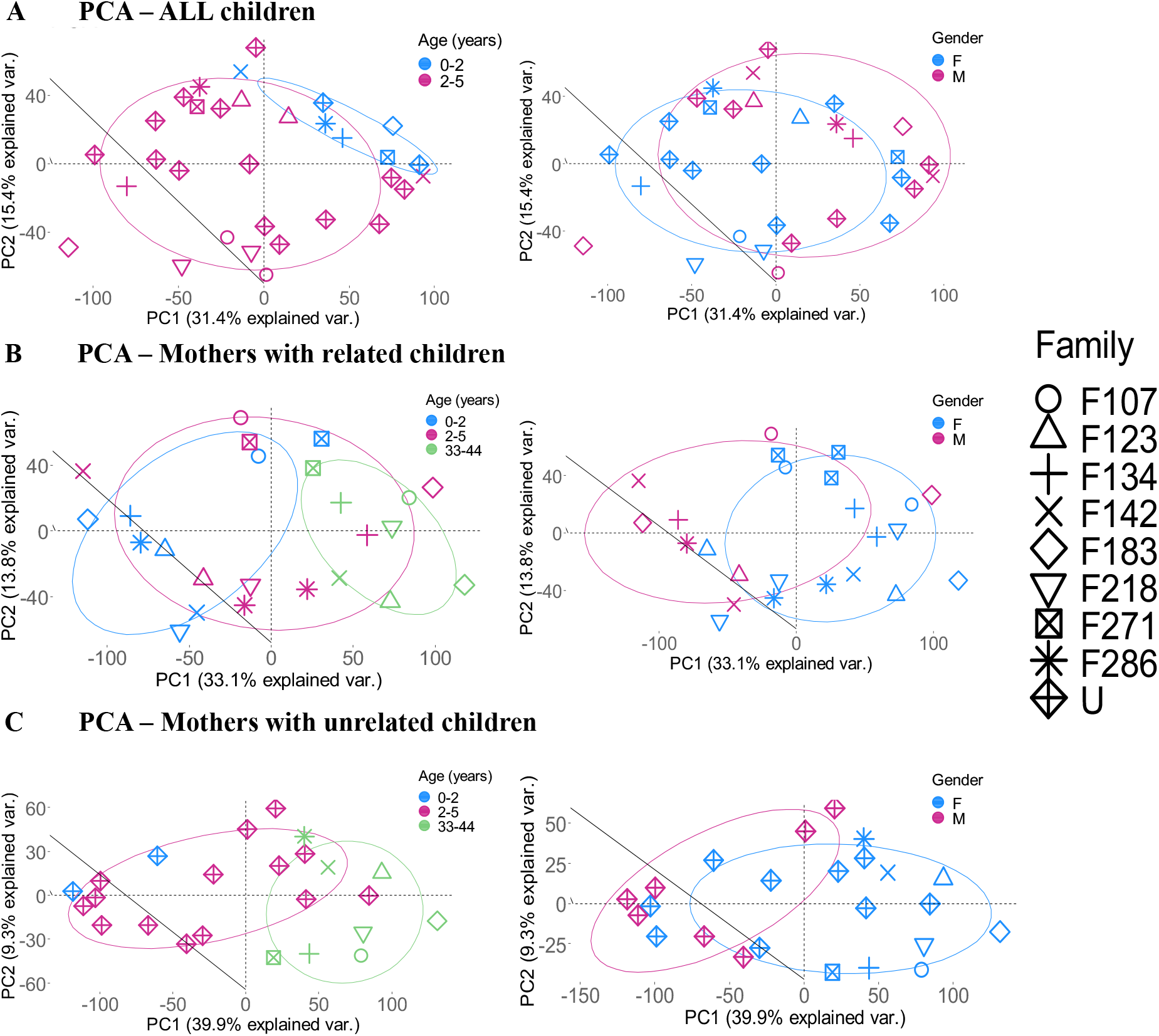
Spatial plots of individuals based on PCA analyses for different population combinations by Age (left side) and Sex (right side). The younger children (aged 0-2) could be separated out from older populations in age-based analyses for all 3 populations.

## Notes

### Competing Interest Statement

The authors have declared no competing interest.

